# Effects of Five Products on Gut Microbiota and Short-Chain Fatty Acids of Obese Rats

**DOI:** 10.1101/652495

**Authors:** Simin Tan, Juan Wang

## Abstract

The effects of banana powder(BP), konjac powder(KP), resistant dextrin(RD), corn starch(CS) and L-carnitine(LC) on gut microbiota and metabolites (SCFA) were evaluated in order to provide basic data for the development of weight-loss functional food. The gut microbiota profile using 16S V4 rDNA high-throughput sequencing technique suggested that the rats of RD, BP and CS group developed an increased richness and diversity in the gut bacterial community, while the abundance of the KP and LC group was not enhanced obviously. *Verrucomicrobiaceae*, *Coprococcus-2* and *Lachnospiraceae* were the main bacterial genera in the CS, BP and RD group respectively, indicating their potential use as prebiotics. On the other hand, rats fed with BP, KP, CS and RD contributed higher total SCFA than those feeding with LC diet. Thus, RD, BP, CS, KP could moduate gut microbiota and increase the SCFA concentration, while the effect of LC are not apparent.

## INTRODUCTION

In recent years, people’s dietary structure has gradually shifted to the consumption of higher quantities of sugars and fats. Long-term intake of high fat can lead to excess accumulated energy in the body as triglycerides, accompanied by abnormal metabolism of sugar, fat, water and salt absolutely. The intestinal environment especially the gut microbiota composition was altered, weakening the gut barrier function and increasing the permeability of the gut, inducing various chronic metabolic disease such as cardiovascular disease, diabetes, fatty liver and obesity. Dietary nutrients can be metabolized by gut microorganisms to improve chronic diseases caused by high-fat diet, which is also a popular research trend in these years^1^. Banana powder (BP), corn starch (CS), konjac powder (KP), resistant dextrin (RD) and L-carnitine (LC) were investigated in this paper, and the changes of gut microorganism composition and metabolites in the obese rats after intervention of five products were mainly discussed to provide academic data for the development of weight-loss functional products.

Banana belongs to the perennial herb of genus Musa, which is rich in protein, carbohydrates, various vitamins and mineral elements that is beneficial to human health. It has been proved to have high nutritional and medicinal value and considered one of the most important economic crops in the south of China^2^. The purity of banana resistant starch extracted from green bananas by low temperature restriction enzyme hydrolysis technique could reach more than 90%^3^. RS has been recommend to be defined as the starch and starch degradation products that can not be digested and absorbed in the small intestine of the human body, but can be fermented by gut microbiota in the large intestine^4^. The fermented end-products of intestinal microorganism are mostly short chain fatty acid, mainly consisted of acetic acid, propionic acid and butyric acid, providing the main energy source of host bacteria. SCFA may play a variety of important roles in metabolism by reducing the pH of gut environment, boosting the probiotics and preventing the pathogenic bacteria^5^.

Recent studies have confirmed that banana powder led to a significant supplement in *Bifidobacteria*, *Bacteriobacteria*, *Lactobacillus* and restriction in *Enterococcus*, given obvious evidence to be associated with increased fermentation products and reduced pH in the gut, thereby improving the condition of intestinal^6^. Moreover, continuous administration of resistant maltodextrin could exert the structure of the cecum microorganism in rats, increasing the *Bacteroidetes* in the cecum and the content of butyrate significantly^7^. Furthermore, intervention of high-amylose corn starch has been demonstrated to adjust colonic flora in the metabolism process, reducing the activity of glucosidase and producing short chain fatty acid especially butyric acid, thus inhibiting the formation of colon cancer in the rats^8^. In addition, the main component of konjac powder is konjac glucomannan (KGM), which also has beneficial effects by selectively promoting the growth of *Anaerobes* and *Lactobacilli* in the intervention group, and reducing the count level of *Faecal Clostridium perfringens* and *Escherichia coli* at the same time^9^. What’s more, L-carnitine is similar to amino acid turning fat into energy. In general, it was identified as the most safe weigh-loss diet supplement by the International Obesity Health Organization in 2003. On the other hand, L-carnitine has negative impact on the cecal microorganism in mice by enhancing the synthesis of trimethylamine (TMA) and trimethylamine-N-oxide (TMAO), increasing occurrence probability of diseases such as atherosclerosis^10^.

Functional modification of banana powder, konjac flour, corn starch, resistant dextrin and L-carnitine have been shown to relieve the levels of weight, blood sugar and blood lipid in high-fat diet fed rats so that they are often used as dietary ingredient for improving health, but the direct cause and effect relationship between the gut microbiota and obesity remains completely unclear. Thus, the reactions of above five products on the intestinal microecology environment, in particular focusing on the intestinal flora and the metabolite (SCFA). Therefore, it is essential to ascertain the potencial protection effects associated with intervention of these products in order to provide basic data for the selection of raw materials and ingredients when developing weight-loss products.

## MATERIALS AND METHODS

### Materials

Banana powder was provided by Tian Jiao Healthy Food (Guangdong) Limited Liability Company. Orlistat was acquired from Chongqing Zhien Pharmaceutical Co., Ltd. Konjac flour, resistant dextrin, corn starch, L-carnitine were all purchased from the market and stored at room temperature following the manufacturer’s instructions.

### Reagent

Standard samples of acetic acid, propionic acid, butyric acid were obtained from Shanghai Yuanmu Biological Technology Co., Ltd. Sodium hydroxide, concentrated hydrochloric acid, metaphosphoric acid, ethyl alcohol, ether, and potassium dihydrogen phosphate were of analytical grade and purchased from the Congyuan Chemical Reagent Co., Ltd.

### Animals and Diets

All the experimental procedures involving animals were approved by Animal Ethics Committee of Animal experiment center of South China University of Agriculture (Guangdong, China, permission number: 2015-B13). After one week of acclimatization, 64 male Sprague-Dawley rats (8 weeks of age, weighing 100±10 g, and received from the medical laboratory animal center of Guangdong Province, SCXK: 2013-0002) were randomly divided into two groups as follows: normal control group(NC) fed with basal diet without intervention (n=8) and obesity model group fed with high-fat diet (n=56). Obesity model was established successfully if the body weight of obesity model group was 20% more over than normal control group(NC). Then, the obese rats were randomly assigned to seven groups: obesity control group (OC), positive control group (PC), banana powder group (BP), konjac powder group (KP), corn starch group (CS), resistant dextrin group (RD) and L-carnitine group (LC). Five groups of products were administered for 6 weeks continuously in accordance with the dosage of 2.5 g/ (kg. BW. D). The NC group and the OC group were given equal volume of distilled water, and the PC group was delivered by gavage to the rats with orlistat. Rats were housed in groups of 4 in each plastic cage with free access to drinking water, under controlled conditions of humidity (40-60%), light(12h/12h light/dark cycle) and temperature (20-25°C). The consumption of food, water and mental state were observed daily. Feces were collected regularly and weight of each rat was measured once a week. At the end of gavage period, rats were sacrificed in a state of anesthesia. The gut tissues were quickly removed and frozen in liquid nitrogen (−80°C) immediately.

### Composition of diet

Ordinary diets and high fat diets were provided by Guangdong medical experimental animal center (SCXK:2013-0002). The nutritive components of high-fat diets were analyzed as follows: 64% sustained feed, 15% lard, 15% sucrose, 5% casein, 0.6% calcium hydrogen phosphate and 0.4% stone powder (processing number: 20170916).

### SCFA measurements in feces

High Performance Liquid Chromatography (HPLC) method was adopted to determine contents of SCFA in the faeces according to Murugesan et al^11^. 6 grams feces of rats was taken in the centrifuge tube and diluted with 20ml deionized water. The contents were centrifuged at 3000 r/min for 10 min. The supernatant was collected and added with 3ml ether and 0.1ml concentrated HCl. The mixture was mixed with a vortex and then centrifuged at 3000 r/min for 10 min again. Then the ether was replaced by 1ml 1mol/L sodium hydroxide and repeated the same steps. The sample was filtered by 0.22 micron micropore membrane at the last. SCFA was quantitatively analyzed using HPLC (Essentia LC-15C, Japan) on a Acclaim 120 C18 liquid chromatography column (4.6mm×250mm×5μm; Agilent) under the following conditions: the flow rate: 1ml/ min, the detection wavelength: 210nm, the column temperature: 30°C, the sampling quantity: 10ul and the flow phase: phosphoric acid buffersolution that the PH was 2.86(A) and acetonitrile(B).

### DNA Extraction, Amplicon Generation, PCR Quantification, Purification and Sequencing

Total genome DNA from samples was extracted using CTAB/SDS method. DNA concentration and purity was monitored on 1% agarose gels. According to the concentration, DNA was diluted to 1ng/μL using sterile water. 16S rRNA/18SrRNA/ITS genes of distinct regions(16SV4/16SV3/16SV3-V4/16SV4-V5, 18S V4/18S V9, ITS1/ITS2, Arc V4) were amplified used specific primer(e.g. 16S V4: 515F-806R, 18S V4: 528F-706R, 18S V9:1380F-1510R, et. al) with the barcode. All PCR reactions were carried out with Phusion®High-Fidelity PCR Master Mix (New England Biolabs). Mix same volume of 1X loading buffer (contained SYB green) with PCR products and operate electrophoresis on 2% agarose gel for detection. Samples with bright main strip between 400-450bp were chosen for further experiments. PCR products was mixed in equidensity ratios. Then, mixture PCR products was purified with Qiagen Gel Extraction Kit(Qiagen, Germany). Sequencing libraries were generated using TruSeq®DNA PCR-Free Sample Preparation Kit(Illumina, USA) following manufacturer’s recommendations and index codes were added. The library quality was assessed on the Qubit@2.0 Fluorometer (Thermo Scientific) and Agilent Bioanalyzer 2100 system. At last, the library was sequenced on an Illumina HiSeq2500 platform and 250 bp paired-end reads were generated.

### Bioinformatic analysis

Paired-end reads was assigned to samples based on their unique barcode and truncated by cutting off the barcode and primer sequence. Paired-end reads were merged using FLASH (V1.2.7, http://ccb.jhu.edu/software/FLASH/)^12^. Sequences with ≥97% similarity were assigned to the same OTU. Representative sequence for each OTU was screened for further annotation. OTU abundance information were normalized using a standard of sequence number corresponding to the sample with the least sequences. Subsequent analysis of alpha diversity and beta diversity were all performed basing on this output normalized data. Alpha diversity is applied in analyzing complexity of species diversity for a sample through6indices, including Observed-species, Chao1, Shannon, Simpson, ACE, Good-coverage. All this indicesin our samples were calculated with QIIME(Version 1.7.0)and displayed with R software(Version 2.15.3). Beta diversity analysis was used to evaluate differences of samples in species complexity. Beta diversity on both weighted and unweighted unifrac were calculated by QIIME software (Version 1.7.0).

### Statistical methods

The data were analyzed with SPSS 21.0 program by one-way ANOVA. The two groups were compared with T test method and Duncan analysis was evaluated for significant difference(P<0.05) in multiple groups.The result was presented as the mean ± standard deviation.

## RESULTS

### OTU and abundance analysis

OTU was used to explain the species richness of the samples preliminarily^13^. The shared and unique OTU among different diet groups is displayed in Fig 1. There were 1209 total OTU and 664 shared OTU in the groups. The unique OTU of RD, BP, CS, KP, LC, NC, OC and PC were 439, 74, 13, 5, 3, 3, 6 and 2 respectively. For the five intervention groups, the highest OTU achieved in the RD group, followed by BP, CS, KP, and lowest in the LC group.

**Fig 1.**
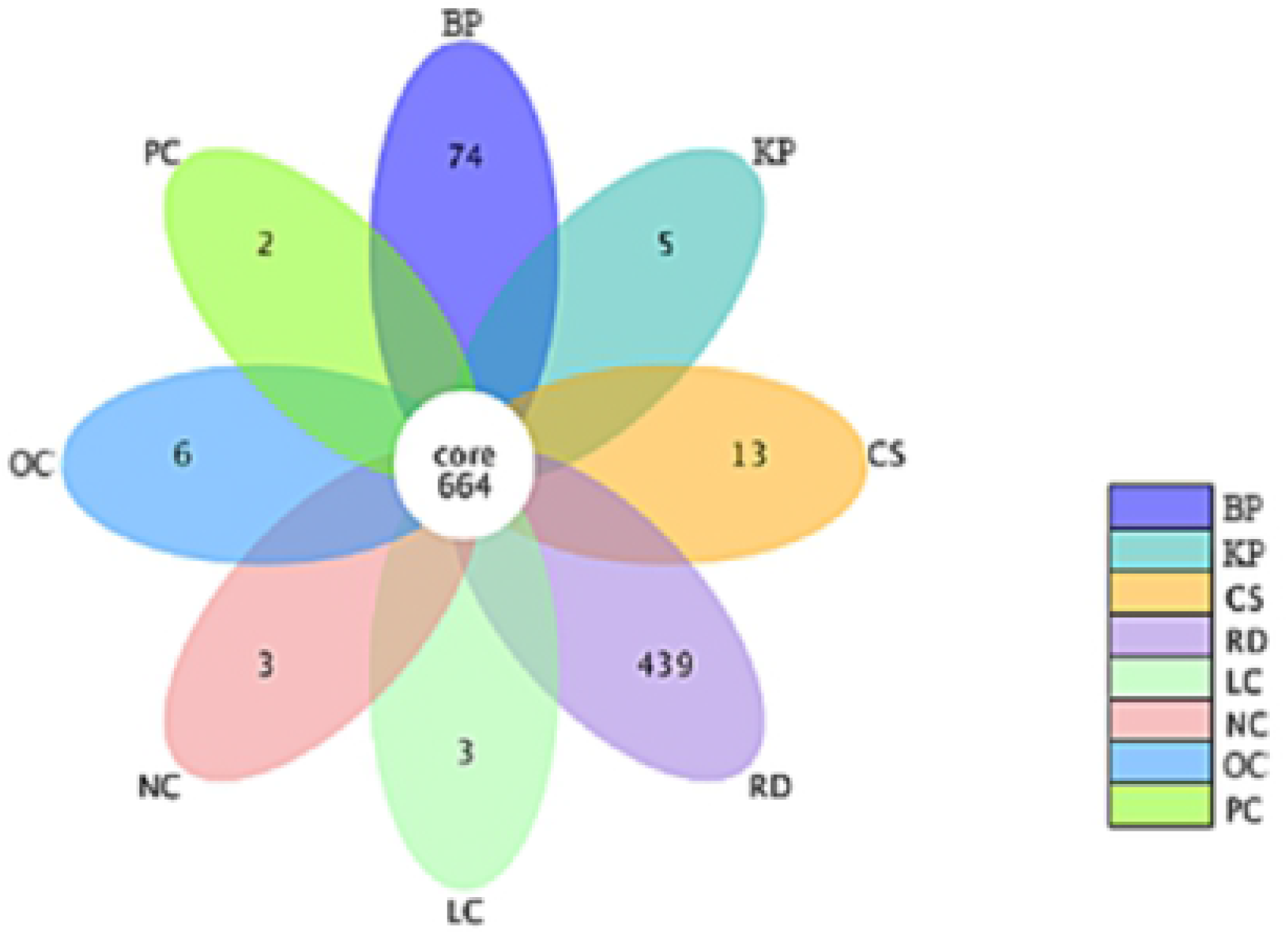
Petal diagram displays numbers of unique and shared OTU among resistant dextrin group(RD), banana powder group(BP), corn starch group(CS), konjac powder group(KP), L-carnitine group(LC), normal control group(NC), obesity control group(OC) and positive control group(PC) (n=8 per group) of rats after 6weeks.

### Alpha diversity analysis

The species diversity of the samples is divided into species abundance and the species uniformity^14^. The Observed species index, the Chao1 index, the Ace index, the Shannon index and the Simpson index are calculated on the basis of OTU and presented in Table 1.The microbial richness can be verified by the Chao1 index and a larger Chao index is associated with a higher community richness. The Chao index applies the following order, from high to low: RD, NC, OC, PC, KP, BP, CS, and LC, indicating that RD supplementation could improve the species richness of intestinal flora in the context of a high-fat diet. Gut microbial uniformity was assessed by the Shannon index. Higher Shannon index is related to increased community diversity.Samples from the NC group showed the highest bacterial community diversity while the lowest uniformity was seen in the CS group.

**Table 1.** Index of alpha diversity in each group

| groups | observed species | chao1 | ace | shannon | simpson |
| --- | --- | --- | --- | --- | --- |
| NC | 778.00 | 824.89 | 825.25 | 7.65 | 0.9903 |
| OC | 788.00 | 824.88 | 837.70 | 7.34 | 0.9870 |
| PC | 783.33 | 822.39 | 825.55 | 7.60 | 0.9893 |
| BP | 754.33 | 804.73 | 803.73 | 7.04 | 0.9837 |
| KP | 768.33 | 818.96 | 821.04 | 7.02 | 0.9787 |
| CS | 730.00 | 803.15 | 798.89 | 6.60 | 0.9403 |
| RD | 760.00 | 830.07 | 800.72 | 7.34 | 0.9847 |
| LC | 756.00 | 796.64 | 802.77 | 7.34 | 0.9877 |

<sup>a</sup>Data are presented as the mean ± SD (n=8).

### Beta diversity analysis

Beta diversity analysis mainly focuses on depicting the variance of different microbial communities among samples. The weighted_UniFrac and unweighted_UniFrac indices were selected and displayed in Fig 2. and Fig 3. respectively. The former index is related to the existence and abundance of species while the latter index only concentrated on the presence and absence of species. The deeper yellow color in the Heatmap means the larger index, representing the larger difference of the species between the samples^15^. If only the presence and absence of the species have been examined (analyzed using the unweighted_unifrac index), the deepest yellow region appeared in the RD group when compared with the other groups, and the difference from high to low was BP > LC > PC > NC > CS > KP > OC (the difference coefficients were 0.617, 0.574, 0.570, 0.567, 0.513, 0.497, 0.492, respectively), implying that the RD group was independent relatively, followed by the BP group. In the five treatment groups, significant alteration of the microbial community only in richness was in accordance with the order: RD > BP > LC > CS > KP (the difference coefficients were 0.492, 0.373, 0.225, 0.164, 0.120 respectively). If the diversity of the species was examined at the same time (analyzed according to the weighted_unifrac index), the deepest yellow region was more concentrated in the CS group when compared with the other groups, followed by the KP group. The order of CS > KP > RD > LC > BP was observed when compared with the OC group (the difference coefficients were 0.113, 0.112, 0.082, 0.08, 0.075, respectively).

**Fig 2.**
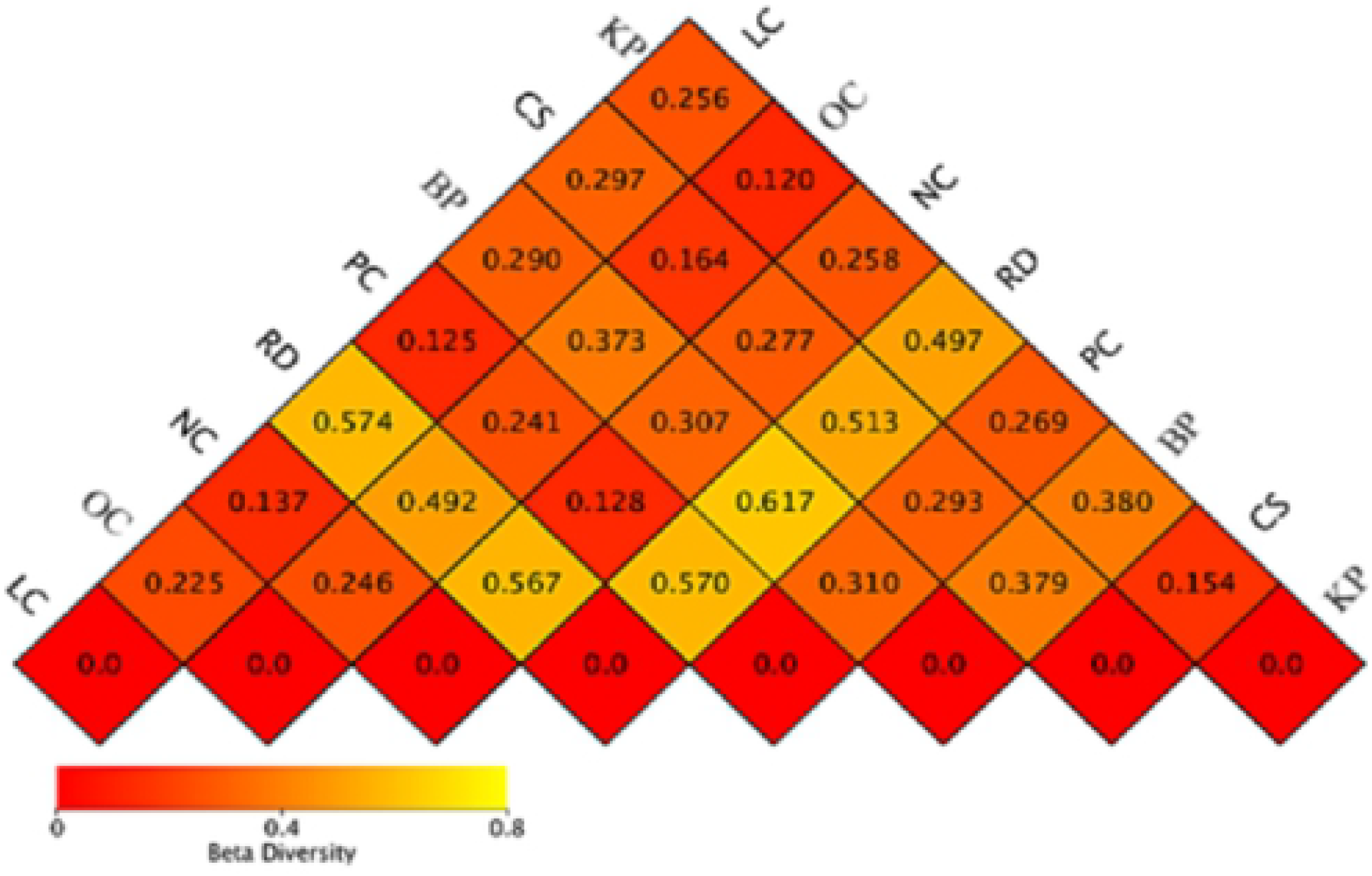
Heatmap based on unweighted_unifrac index

**Fig 3.**
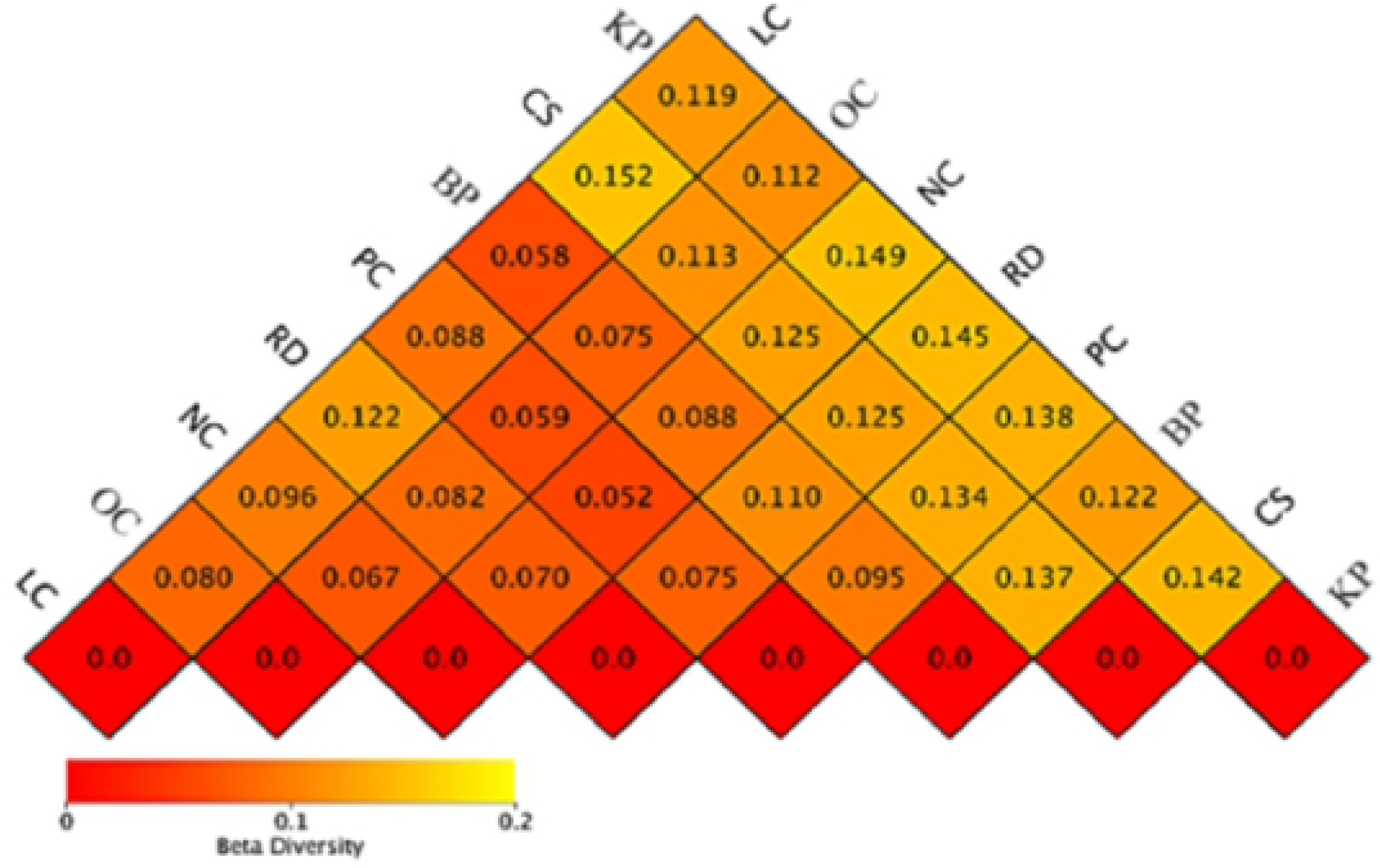
Heatmap based on weighted_unifrac index

### UPGMA cluster analysis

The sample also can be cluster analyzed based on the unweighted_UniFrac index and the weighted_UniFrac index which exhibited in Fig 4. and Fig 5. respectively. The system evolution tree of the sample was set up to further investigate the similarity of the species composition in the samples, in which the closer sample distance and the shorter branch length suggests the more similarity of the species composition between samples^16^. The RD group was viewed as the most independent sample based on the analysis of the unweighted_UniFrac index, followed by the BP group, while the CS group is the most independent if the weighted_UniFrac index considered and followed by the KP group. This analysis was consistent with the above Beta diversity Heatmap analysis, which supply further evidence that results of microbial community depending on two exponents also appeared difference.

**Fig 4.**
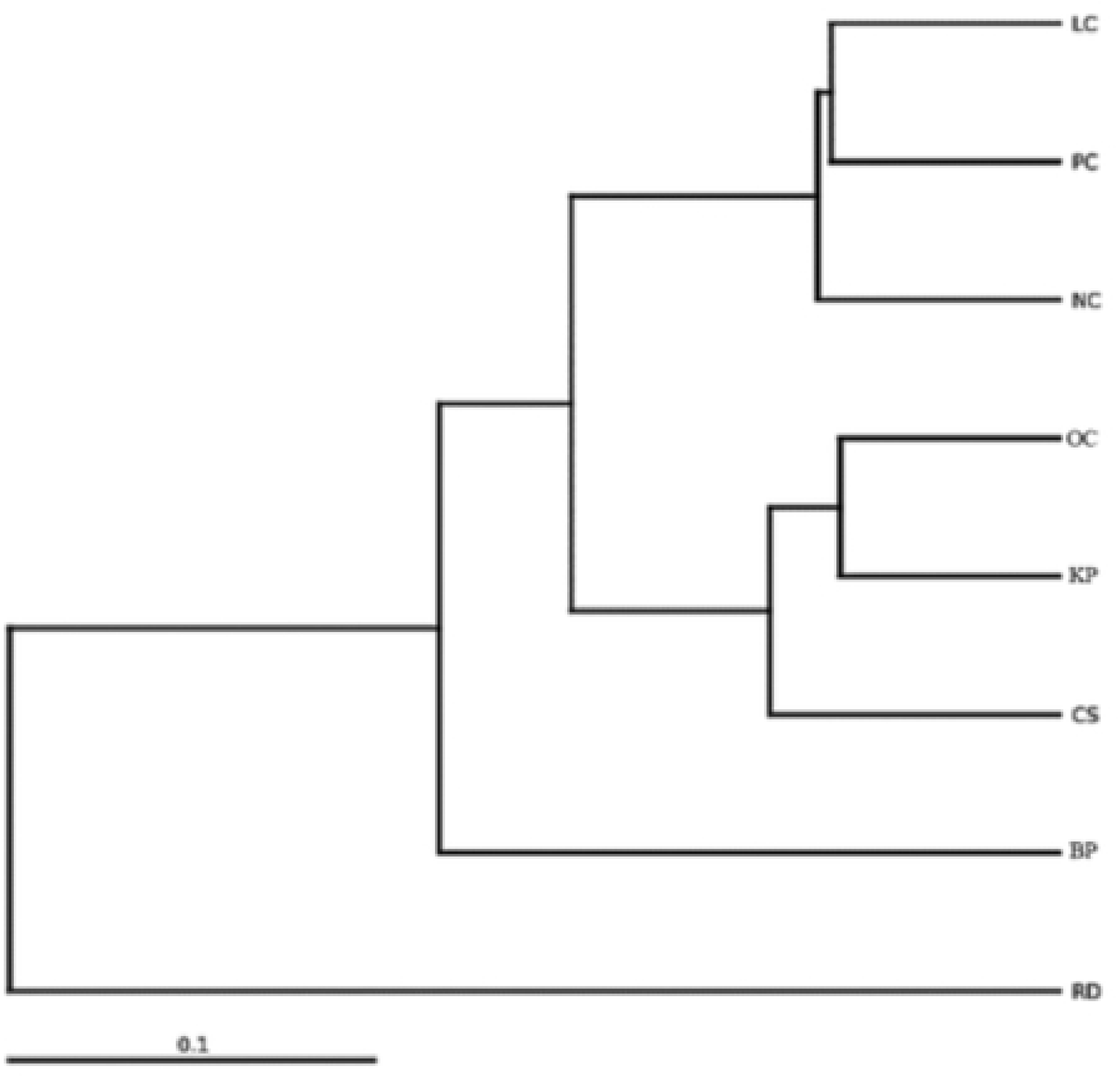
Cluster analysis diagram based on unweighted UniFrac index

**Fig 5.**
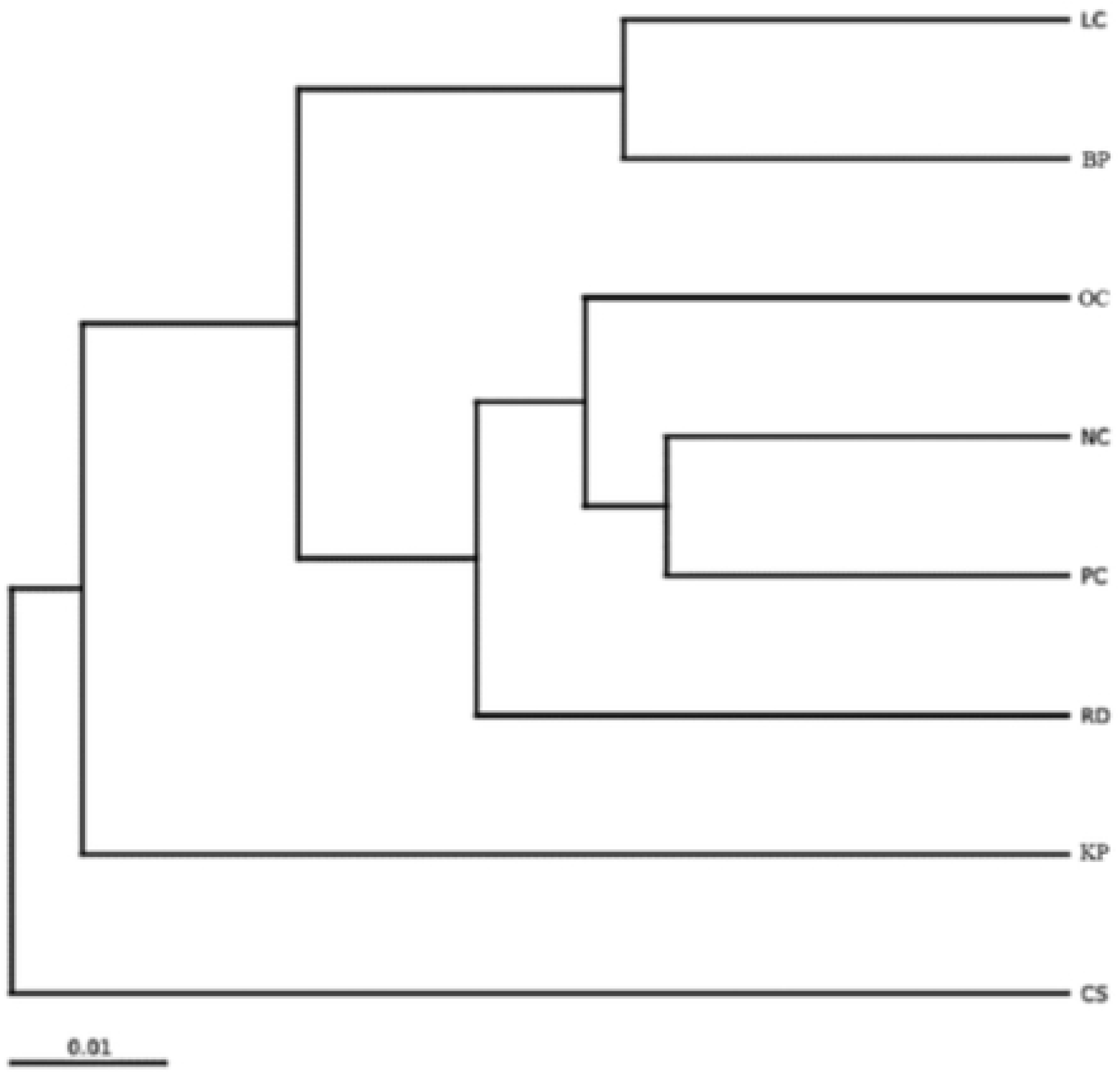
Cluster analysis diagram based on weighted UniFrac index

### Analysis of gut microbiota at the phylum level

Influences of the gut microbiota structure in five administration groups was as reflected in Fig 6.The three dominant Phylum in all the groups were Firmicutes, Bacteroidetes and Proteobacteria, accounting for 65.2%, 24.1% and 5.6% of the total eumycota, respectively. The abundance of Proteobacteria in KP group (13.3%) was significantly higher than that of other groups (abundance <7%), and the relative abundance of Verrucomicrobia in CS group was the highest (12.8%), while less than 2% was detected in other groups. More importantly, the proportion of Bacteroidetes and Firmicutes was the highest in the BP group (50.8%) and lowest in the OC group(29.2%).

**Fig 6.**
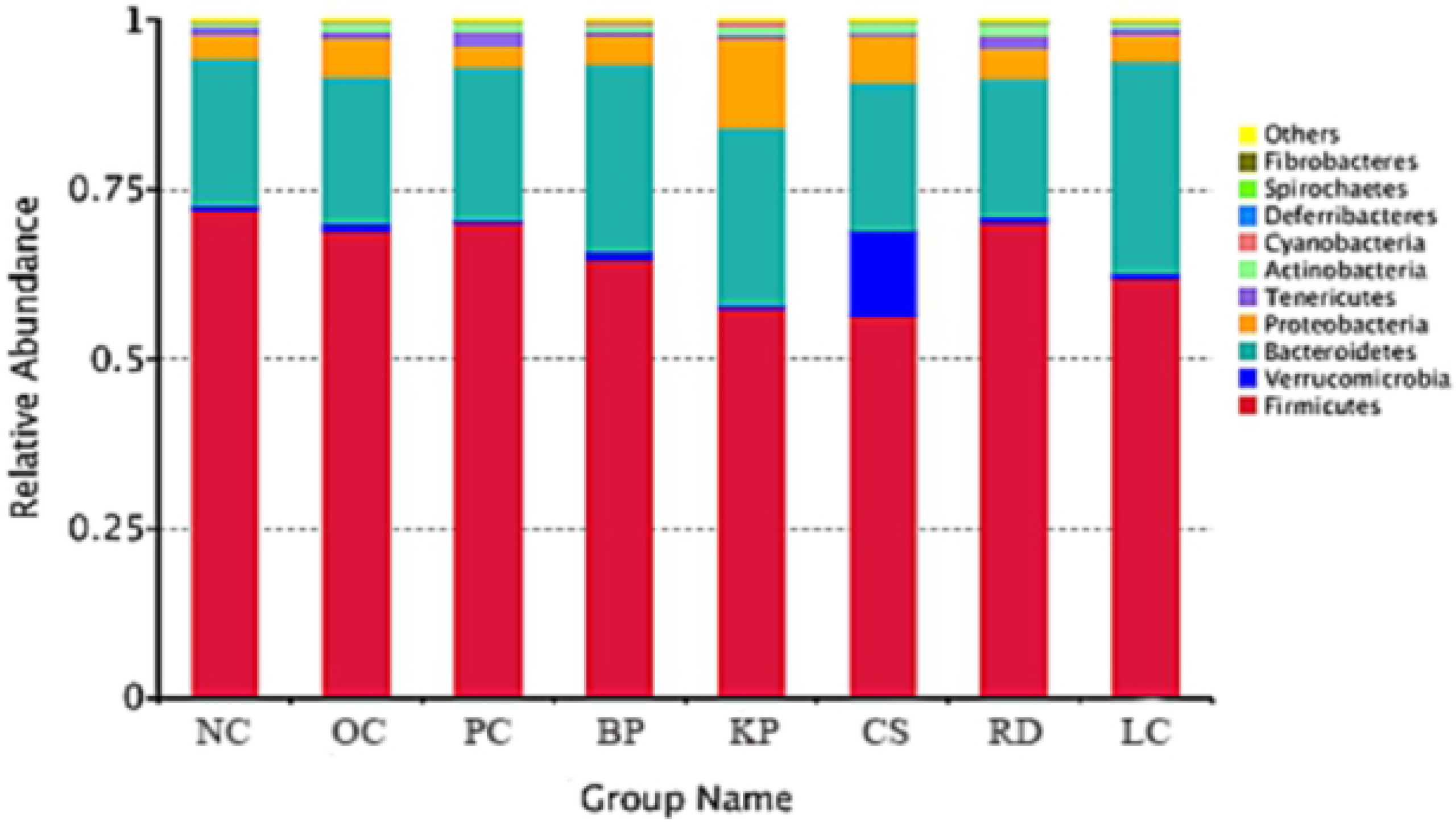
Composition of gut microbiota in each group at the phylum level

### Analysis of gut bacterial population at the genus level

Comparisons of gut microbiota at the genus level are shown in Fig 7. The three genera with the highest relative abundance are: *Lachnospiraceae_NK4A136_group* (7%), *Bacteroides* (6%), and *Ruminococcaceae* (7%). The abundance of *Coprococcus-2* (4%) in the BP group was significantly higher than that in the other groups, and the supplemention of orlistat significantly enhanced the growth of *Ruminococcaceae_NK4A214_group*(6%). Furthermore, the most positive environment of *Akkermansia* existed in the CS group(13%), and the highest abundance of the genus *Lachnospiraceae_NK4A136_group* was revealed in the RD group(9%). The *Coprococcus-2*, *Ruminococcaceae_NK4A214_group*, *Akkermansia* and *Lachnospiraceae_NK4A136_group* all belong to beneficial bacteria. In addition, compared with the OC group, the abundances of *Bacteroides*, *Turicibacter*, *Olligella*, *Desulfovibrio* and *Romboutsia* of five products follow that order: RD < CS < BP < KP < LC. However, the above five generas were distributed the highest in the LC group, taking the proportions of 10%, 4%, 3%, 7% and 4% respectively, and the contents of beneficial bacterial genera such as *Ruminococcaceae*, the *Lachnospiraceae_NK4A136_group* and the *Akkermansia* were low in LC, KP and OC groups.

**Fig 7.**
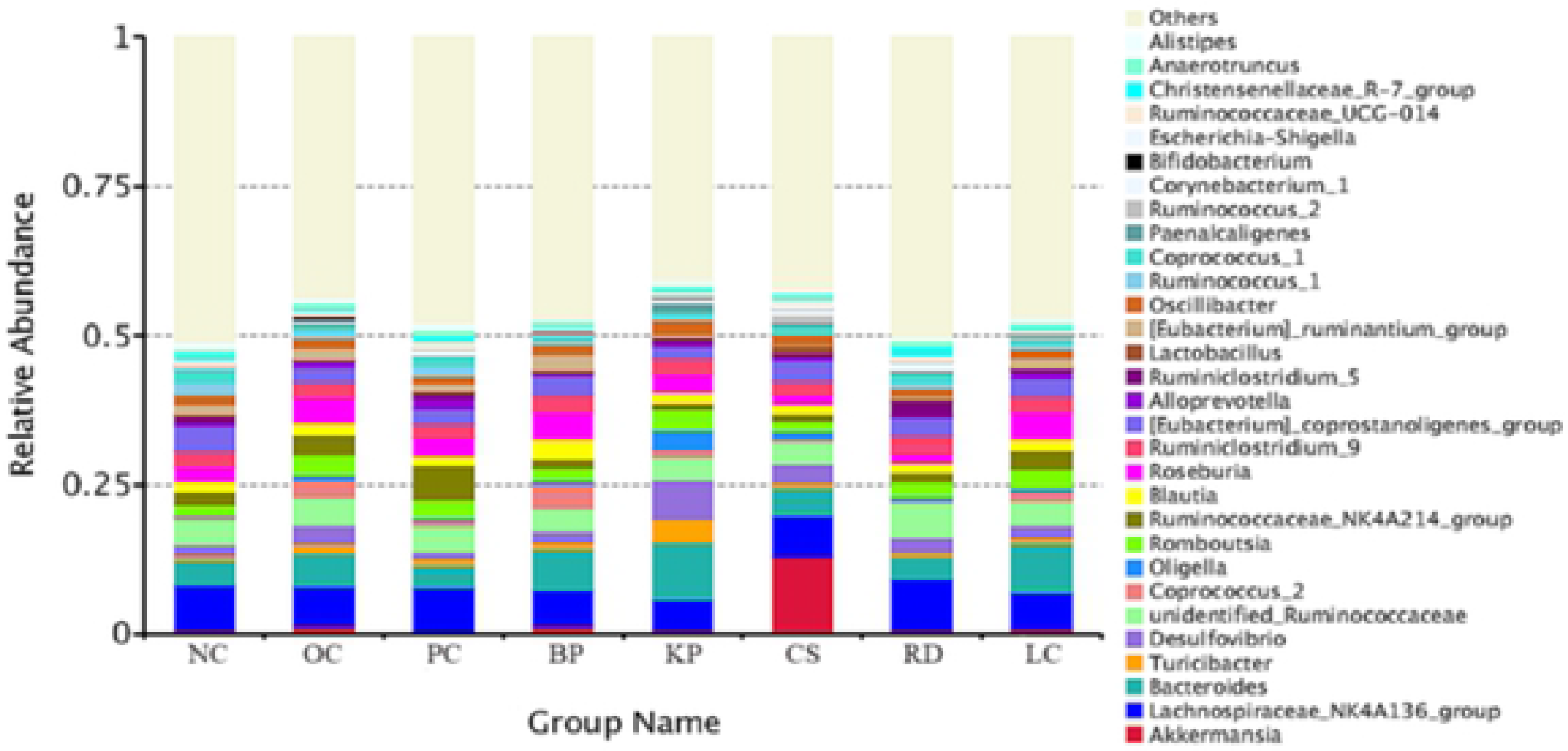
Composition of gut microbiota in each group at the genus level

### Analysis of the fecal SCFA content

The different effects of consumption of the five products on the production of the major SCFAs (acetate, propionate, butyrate) in the gut are summarized in Table 2. The total SCFA content in the feces of the RD group was the highest compared with the NC group (P<0.05), while it was decreased significantly in the LC and OC group(P<0.05). Similar increased content of SCFAs were found in the BP, KP, CS and PC group, but there was no statistical difference with the NC group. Compared with the OC group, SCFAs in the PC, BP, KP, CS and RD group increased significantly (P<0.05), indicating that BP, KP, CS and RD could promote the production of SCFA, among of which the effect of RD was the best, while LC had no obvious effect. The content of acetic acid in BP, KP, CS and RD group was significantly higher than that of the OC group (P<0.05), while it was significantly lower than that of the NC group in the LC group (P<0.05). The RD group and LC group achieved higher content of propionic acid than the OC group significantly (P<0.05), while rats fed with the orlistat, BP, KP and CS has lower content of propionic acid than that of the NC group (P<0.05). The content of butyric acid in the RD group was the highest, and there was significant difference with the OC group (P<0.05). The content of propionic acid also increased in BP, KP, CS and LC group, but there was no statistical difference compared with the NC or OC group.

**Table 2.** Fecal content of SCFA in the control and treated group(μmol/ml)

| groups | NC | OC | PC | BP | KP | CS | RD | LC |
| --- | --- | --- | --- | --- | --- | --- | --- | --- |
| acetate | 5.29<sup>#</sup>±0.26 | 2.18<sup>*</sup>±0.08 | 2.48<sup>*</sup>±0.02 | 5.31<sup>#</sup>±0.59 | 4.81<sup>#</sup>±0.49 | 5.24<sup>#</sup>±0.41 | 5.53<sup>#</sup>±0.44 | 1.92<sup>*</sup>±0.06 |
| propionate | 2.96<sup>#</sup>±0.06 | 1.85<sup>*</sup>±0.32 | 1.49<sup>*</sup>±0.33 | 1.30<sup>#*</sup>±0.05 | 1.80<sup>*</sup>±0.03 | 1.45<sup>*</sup>±0.04 | 2.27<sup>#</sup>±0.13 | 2.22<sup>#</sup>±0.26 |
| butyrate | 2.32±0.30 | 1.39±0.14 | 1.40±0.21 | 2.20±0.14 | 2.15±0.27 | 2.74±0.44 | 2.86<sup>#</sup>±0.57 | 2.75±0.28 |
| SCFA | 9.47<sup>#</sup>±0.07 | 6.53<sup>*</sup>±0.37 | 8.38<sup>#</sup>±0.10 | 8.81<sup>#</sup>±0.63 | 8.76<sup>#</sup>±0.21 | 9.42<sup>#</sup>±0.67 | 10.66<sup>#*</sup>±0.28 | 6.89<sup>*</sup>±0.30 |

<sup>a</sup>Data are expressed as the mean ± SD (n=8). <sup>#</sup>Different lowercase symbols above the same column indicate a significant difference compared with NC and OC group respectively (P < 0.05).

## DISCUSSION

Effects of five kinds of products including banana powder(BP), konjac powder(KP), corn starch(CS), resistant dextrin(RD) and L-carnitine(LC) to regulate the intestinal microorganism composition and metabolite SCFAs of high-fat diet rats have been studied. The results of OTUs showed that RD, BP and CS could enhance the abundance of intestinal flora in obese rats compared with the OC group, while the effect of KP and LC was not apparent. At the level of genus, compared with the OC group, in the terms of growth of beneficial bacteria, the corn starch promoted the growth of *Verrucomicrobiaceae* significantly. The content of the *Coprococcus-2* and *Lachnospiraceae* was the highest in the BP and RD group respectively, reflecting the positive effect of CS, BP, and RD to proliferate the beneficial bacteria. Compared with the OC group, the inhibitory effect of RD on *Bacteroides*, *Turicibacter*, *Olligella*, *Desulfovibrio* and *Romboutsia* was the best, followed by CS, BP, KP and LC. The highest content of SCFAs in each group was acetic acid and the SCFAs in the four products was significantly higher than that of the obese control group (P<0.05). The content of SCFAs follow the order from high to low: RD, CS, BP and KP group, while L-carnitine group was the lowest.

Beta diversity analysis revealed that the RD was a relatively independent sample according to the unweighted_unifrac index, and the OTUs results also showed that the abundance of intestinal flora was the highest in the RD group. The total SCFAs content and the content of butyric acid in the feces of the RD group were the highest, indicating that the species richness of RD was improved more greatly and effectively than that of other four products, which can promote the production of more SCFAs by metabolic process of microorganisms and improve intestinal health. The structure of intestinal flora in rats was further analyzed at the genus level. *Lachnospiraceae_NK4A136_group* was found as the predominant genus of RD, which is a weakly gram-positive bacteria and can ferment glucose to produce formic acid, lactic acid, acetic acid, etc., but succinic acid, butyric acid and propionic acid can not be produced. Li ewska K (2015) report that it had a positive effect on the composition of intestinal flora in rats fed with RD. The count of *Bifidobacterium* and *Lactobacillus* in the feces and cecum increased, and the count of *Clostridium* and *Bacteroides* decreased.The stimulation of *Bifidobacterium*, *Lactobacilli* and SCFA may help inhibit the pathogenic microorganisms in the colon^17^. Therefore, the higher butyric acid content in the RD group may be produced by *Bifidobacterium* and *Lactobacillus* in this research.

As the type of RS2 resistant starch, banana powder(BP) and corn starch(CS) can play a good regulatory role in the intestinal flora of obese rats. J Yang (2013) found that both RS1 and RS2 can promote growth of *Bifidobacterium*. The count of lactobacilli, Streptococcus and Enterobacteriaceae in the cecum of rats in RS2 diets was significantly higher than that in the rats with RS1 diets (P < 0.05). The concentrations of propionic acid and propionate in feces and cecum fed with RS2 were significantly higher than those in RS1 group (P < 0.05)^18^. The effect of R2 resistant starch in different sources may be different. In this study, the total SCFAs content and butyric acid content of BP and CS group were high, but the number of OTUs in BP group was higher than the CS group. At the phylum level, the proportion of Bacteroidetes and Firmicutes in the BP group was the highest, and lowest in the OC group. Ley, R. E. (2006) reported that the proportion of Bacteroidetes to Firmicutes in obese individuals was lower than that in healthy individuals^19^. Martinez, I et al. (2010) found that resistant starch can increase the number of Actinobacteria and Bacteroidetes, and reduce the number of Firmicutes in human body^20^. This is consistent with the our research findings in this paper, indicating that the effect of banana powder on the regulation of intestinal microflora may be better than that of corn starch. At the level of the genus, the change of gut microbiota in the two groups was analyzed. *Akkermansia* and *Coprococcus-2* was found as the dominant genus in the BP and CS group respectively. *Akkermansia* is mainly related to the human body weight, the inflammatory state and the glucose tolerance in the cecum and colon digestion. *Coprococcus-2* are common in the gut and feces. Their fermentation products include butyric acid, acetic acid and gas, which can regulate the health of the intestine. In general, in terms of adjusting the structure of intestinal flora, banana powder is better than corn starch.

The effect of KP on the intestinal flora structure and metabolites of obese rats was less reported. AA Elamir et al.(2008) reported that konjac glucomanic hydrolysate (GMH) significantly promoted the growth of *Anaerobes* and *Lactobacilli* in the intervention group (P<0.001), and significantly reduced the count level of *Faecal Clostridium perfringens* and *Escherichia coli* (P<0.001). In this study, the effect of KP on increasing the abundance of intestinal flora was not obvious. In the Beta analysis of unweighted UniFrac index, the difference coefficient of the KP group and the OC group was the lowest. The OTUs results also showed that the abundance of intestinal flora in the KP group was low, but the total fecal SCFAs content in the KP group was similar to that of the CS and the BP group, showing that the intestinal microflora of the KP group had apparent effect on the production of SCFAs, but had no significant effect on the increase of the abundance of intestinal flora. In addition, the abundance of *Bacteroides*, *Turicibacter*, *Olligella*, *Desulfovibrio* and *Romboutsia* in the KP group was relatively high. Z Wang et al. (2018) reported the adverse effects of *Romboutsia* and *Turicibacter*^21^. Thus, the above five species were speculated to have the inhibitory effects on enhancing the abundance of the beneficial genera, which lead to the lower integral abundance of intestinal flora in the KP group.

Studies of Kuka et al. (2014) showed that after the intake of L-carnitine, the intestinal microorganism group can produce trimethylamine (TMA) and organic substance (TMAO), increasing the incidence probability of obesity, cardiovascular and other diseases^22^. The results of this study showed that the abundances of intestinal flora and SCFAs in the LC group were the lowest, which may be related to the above mechanism. On the other hand, the abundances of *Bacteroides*, *Turicibacter*, *Olligella*, *Desulfovibrio* and *Romboutsia* were also the highest in the LC group, which could used to provide a new direction for evaluating the side effects of L-carnitine.

## ACCOCIATED CONTENT

### Supporting Information

The intestinal microbial of colonies of 8 groups of rats were measured. Each group had 3 parallel samples, totaling 24 samples. The detailed statistical results of each sample were shown in Table 1 and Table 2. The percentage of sequence error in each data during filtering step was low relatively. The number of effective bases ranged from 14306945 ± 1184436 to 18986644 ± 1164404, indicating the data was reliable. Tags are formed by overlap between base fragments as shown in Table 2. The number of clean Tags filtered from the raw data was between 71911 ± 6572 and 91461 ± 3768 and the percentage of effective splicing tags was more than 75%, reflecting the effective utilization rate of data was high.

Species dilution curve can directly reflect the rationality of the amount of sequencing data, and indirectly explain the species richness in the sample. When the curve is flat, it shows that the amount of sequencing data is gradually reasonable, representing more data will only produce a few number of new species (OTU). The species number saturation curve (Rank Abundance curve) can directly reflect the species richness and uniformity in the sample. In the horizontal direction, species richness is reflected by the width of the curve. The higher the species richness, the larger the span of the curve on the horizontal axis. In the vertical direction, the smoothness of the curve reflects the homogeneity of species in the sample. The smoother the curve, the more homogeneous the species distribution.

As shown in Fig. 1 and Fig. 2, the species dilution curves of 24 samples tend to be flat with the increase of the amount of sequencing data, indicating that the amount of sequencing data was reasonable and the data reliability was high. According to the Rank Abundance curve, the span of RD group was the largest (about 0-1200) in the abscissa coordinates, and the span of the other groups was similar (about 0-800), showing that the richness of intestinal microorganisms in the RD group was the highest, while the richness of other groups was slightly different. In the vertical direction, it can be seen that the tail end of each group of curves appeared obvious convex folds, suggesting that the overall homogeneity of each group of samples was low.

## AUTHOR INFORMATION

### Notes

The authors declare no conflict of interest.

## REFERENCES

1. X He, W Sun, T Ge et al.: An increase in corn resistant starch decreases protein fermentation and modulates gut microbiota during in vitro cultivation of pig large intestinal inocula. Animal Nutrition 2017;3:219–224.

2. Wu Jf, Wang Hm.: Development status and competitiveness analysis of banana industry in China. Modern Agricultural Science and Technology 2013;7:328–330.

3. Wang J, Tang X J, Chen P S, et al.: Changes in resistant starch from two banana cultivars during postharvest storage. Food Chemistry 2014;156:319–325.

4. Englyst H N, Kingman S M, Cummings J H.: Classification and measurement of nutritionally important starch fractions. European Journal of Clinical Nutrition 1992;75:327–338.

5. Bilal Ahmad Ashwar, Adil Gani, et al.: Preparation, health benefits and applications of resistant starch-a review. Starch 2016;68:287–301.

6. Bai Yl, Peng Zf, Chen Qf, et al.: Study on the effect of banana powder intervention on intestinal microflora in diabetic rats. Modern Food Science and Technology 2013;29:2110–2114.

7. S Miyazato, Y Kishimoto, et al.: Continuous intake of resistant maltodextrin enhanced intestinal immune response through changes in the intestinal environment in mice. Bioscience of Microbiota Food & Health 2016; 35:1.

8. S Nakanishi, K Kataoka, et al.: Effects of high amylose maize starch and Clostridium butyricum on microbiota and formation metabolism in colonic of azoxymethane-induced aberrant crypt foci in the rat colon.Microbiology & Immunology 2003;47:951–958.

9. AA Elamir, RF Tester, et al.: Effects of konjac glucomannan hydrolysates on the gut microflora of mice.Nutrition & Food Science 2008;38:422–429.

10. RA Koeth, Z Wang, et al.: Intestinal microbiota metabolism of L-carnitine, a nutrient in red meat, promotes atherosclerosis.Nature Medicine 2013;19:576–585.

11. Murugesan S, Ulloa, Martinez M, Martinez, Rojano H, et al.: Study of the diversity and short-chain fatty acids production by thebacterial community in overweight and obese Mexican children. European Journal of Clinical Microbiology and Infectious Diseases 2015;34:1337–1346.

12. MagočT, Salzberg S L. FLASH: fast length adjustment of short reads to improve genome assemblies. Bioinformatics 2011; 27: 2957–2963.

13. Lundberg, Derek S., et al.: Practical innovations for high-throughput amplicon sequencing. Nature methods 2013;10:999–1002.

14. C Lozupone, M Hamady, and R Knight.: UniFrac-An online tool for comparing microbial community diversity in a phylogenetic context. BMC Bioinformatics 2006;7:371.

15. Avershina, Ekaterina, Trine Frisli, and Knut Rudi.: De novo Semi-alignment of 16S rRNA Gene Sequences for Deep Phylogenetic Characterization of Next Generation Sequencing Data. Microbes and Environments 2013; 28:211–216.

16. Bokulich, Nicholas A., et al.: Quality-filtering vastly improves diversity estimates from Illumina amplicon sequencing. Nature methods 2013;10:57–U11.

17. Śliżewska K, Z Libudzisz, et al.: Dietary resistant dextrins positively modulate fecal and cecal microbiota composition in young rats. Acta Biochimica Polonica 2015; 62:677.

18. J Yang, I Martínez, et al.: Invitro characterization of the impact of selected dietary fibers on fecal microbiota composition and short chain fatty acid production. Anaerobe 2013; 23:74–81.

19. Ley RE, Turnbaugh PJ, Klein S, Gordon JI.: Microbial ecology-Human gut microbes associated with obesity. Nature 2006;444:1022–1023.

20. Martinez I, Kim J, Duffy PR, Schlegel VL, Walter J.: Resistant starches types 2 and 4 have differential effects on the composition of the fecal microbiota in human subjects. PLoS One 2010, 5:e15046.

21. Z Wang, W Zhang, et al.: Influence of Bactrian camel milk on the gut microbiota. Journal of Dairy Science 2018; 101:5758–5769.

22. Kuka J, Liepinsh E, et al.: Suppression of intestinal microbiota-dependent production of pro-atherogenic trimethylamine N-oxide by shifting L-carnitine microbial degradation. Life Sciences 2014;117:84–92.

23. Wang Q, et al.: Naive Bayesian classifier for rapid assignment of rRNA sequences into the new bacterial taxonomy. Applied and environmental microbiology 2007;73:5261.

24. White, James Robert, Niranjan Nagarajan, and Mihai Pop.: Statistical methods for detecting differentially abundant features in clinical metagenomic samples. PLoS computational biology 2009;5:e1000352.

25. Youssef, Noha, et al.: Comparison of species richness estimates obtained using nearly complete fragments and simulated pyrosequencing-generated fragments in 16S rRNA gene-based environmental surveys. Applied and environmental microbiology 2009;75:5227.

26. Lozupone, Catherine A., et al.: Quantitative and qualitative β diversity measures lead to different insights into factors that structure microbial communities. Applied and environmental microbiology 2007;73:1576–85.

